# Lack of authentic atrial fibrillation in commonly used murine atrial fibrillation models

**DOI:** 10.1101/2021.08.17.456655

**Authors:** Fumin Fu, Michael Pietropaolo, Lei Cui, Shilpa Pandit, Weiyan Li, Oleg Tarnavski, Suraj S. Shetty, Jing Liu, Jennifer M. Lussier, Yutaka Murakami, Prabhjit K. Grewal, Galina Deyneko, Gordon M. Turner, Andrew K.P. Taggart, M. Gerard Waters, Shaun Coughlin, Yuichiro Adachi

**Affiliations:** Cardiovascular & Metabolic Diseases, Novartis Institutes for BioMedical Research, Inc. Cambridge, Massachusetts, United State of America

## Abstract

The mouse is a useful preclinical species for evaluating disease etiology due to the availability of a wide variety of genetically modified strains and the ability to perform disease-modifying manipulations. In order to better characterize atrial fibrillation (AF), we profiled several commonly used murine AF models. We initially evaluated a pharmacological model of acute carbachol (CCh) treatment plus atrial burst pacing in C57BL/6 mice. In an effort to observe micro-reentrant circuits indicative of authentic AF, we employed optical mapping imaging in isolated mouse hearts. While CCh reduced atrial refractoriness and increased atrial tachyarrhythmia vulnerability, the left atrial (LA) excitation patterns were rather regular without reentrant circuits or wavelets. Therefore, the atrial tachyarrhythmia resembled high frequency atrial flutter, not typical AF *per se*. We next examined both a chronic angiotensin II (Ang II) infusion model and the surgical model of transverse aortic constriction (TAC), which have both been reported to induce atrial and ventricular structural changes that serve as a substrates for micro-reentrant AF. Although we observed some extent of atrial remodeling such as fibrosis or enlarged LA diameter, burst pacing-induced atrial tachyarrhythmia vulnerability did not differ from control mice in either model. This again suggested that an AF-like pathophysiology is difficult to demonstrate in the mouse. To continue searching for a valid murine AF model, we studied mice with a cardiac-specific deficiency (KO) in liver kinase B1 (Cardiac-LKB1), which has been reported to exhibit spontaneous AF. Indeed, the electrocardiograms (ECG) of conscious Cardiac-LKB1 KO mice exhibited no P waves and had irregular RR intervals, which are characteristics of AF. Histological evaluation of Cardiac-LKB1 KO mice revealed dilated and fibrotic atria, again consistent with AF. However, atrial electrograms and optical mapping revealed that electrical activity was limited to the sino-atrial node area with no electrical conduction into the atrial myocardium beyond. Thus, Cardiac-LKB KO mice have severe atrial myopathy or atrial standstill, but not AF. In summary, the atrial tachyarrhythmias we observed in the four murine models were distinct from typical human AF, which often exhibits micro- or macro-reentrant atrial circuits. Our results suggest that the four murine AF models we examined are poor representations of human AF, and raise a cautionary note about the use of any murine model to study AF.

## Introduction

Atrial fibrillation (AF) is the most common cardiac arrhythmia, with prevalence increasing with age [1]. AF can occur without any signs or symptoms, and if left untreated can lead to serious life-threatening complications including heart failure and stroke [2]. Although pharmacologic and non-pharmacologic treatment options for AF are available [3], these approaches have serious limitations. Pharmacologic treatment usually increases atrial refractoriness to inhibit sustained reentry. However, pharmacologic treatments often have ventricular pro-arrhythmic effects, which are contraindicated in some patient populations such as those with AF with concurrent structural heart diseases. Non-pharmacologic treatments such as catheter ablation have issues of surgery complications and AF recurrence. Thus, an unmet medical need for a novel and safe anti-arrhythmic drug for AF remains [1]. To meet this need, establishing a human-translatable animal model is essential for evaluating new anti-arrhythmic therapies. The most commonly used models to date utilize large animals such as dogs, pigs and sheep that develop AF upon rapid atrial pacing [4]. However, due to limited accessibility and high cost of the large animal models, there is a desire to use rodent models.

In recent years, several murine models have been reported to mimic human AF. Among them, a commonly-used model employs acute cholinergic stimulation by carbachol (CCh) treatment followed by burst pacing [5]. CCh treatment mimics parasympathetic stimulation by activating G protein-coupled potassium (GIRK) channels to reduce atrial refractoriness [6], which in turn makes atria vulnerable to re-entry arrhythmias triggered by burst pacing. Two other murine AF models involve production of heart dysfunction by transverse aortic constriction (TAC) surgery or chronic angiotensin (Ang) II-infusion [7-14]. These models exhibit structural and electrical remodeling in atria, and have been reported to be vulnerable to burst pacing-induced AF.

Several genetically modified rodent models have also been reported to have AF [15-20]. Liver kinase B1 (LKB1) is a serine/threonine protein kinase that controls the activity of AMP-activated protein kinase (AMPK) family members. AMPK plays a role in various physiological processes such as cell metabolism, cell polarity, apoptosis and DNA damage response. In the heart, the LKB1-AMPK pathway influences ion channel remodeling, fibrosis induction and apoptosis [21, 22]. Relevant to this work, the Cardiac-specific LKB1 deficient (KO) mouse model has been reported to have persistent AF [23-25] that is evident even without burst pacing.

Here we profiled the above murine models to determine whether they are reasonable representations of human AF. In particular, we evaluated atrial electrical activity during tachyarrhythmia and compared the general characteristics of murine AF models to those reported in human and/or large animal models of AF featuring wavelets and/or micro-reentry [26].

## Material and Methods

All animal studies were approved by the Novartis Institutional Animal Care and Use Committee (IACUC). Institutional protocol numbers used in the studies were 17CVM012, 18CVM031, 18CVM032 and 20CVM023.

### *In vivo* electrophysiology

A 1.1 French octapolar electrode (Transonic, NY, USA) was inserted via the right jugular vein into the right atrium in isoflurane-anesthetized mice. Right atrial (RA) electrograms (EG) and ECG lead II signals were acquired and analyzed using a Power lab system (ADinstruments, Colorado, USA). For CCh studies, CCh (Tokyo Chemical Industry, Catalog # C0596, 0.3 mg/kg in saline) was injected intraperitoneally in male 9-11 week old C57BL/6J mice. To evaluate atrial effective refractory period (ERP) and AF inducibility, electrical stimulations were sent to the octapolar electrode using a programmable stimulus generator (STG4004, Multichannel systems, Reutlingen, Germany). For AF inducibility, 5 seconds of electrical burst pacing stimulations with 33, 50 and then 100 Hz rectangular pulses (2 ms width) were applied 6 times at each frequency (total 18 burst pacings). More than 0.5 s of continuous high frequency and polymorphic atrial EG signal after termination of burst pacing was counted as a positive tachyarrhythmia. In most of the cases, tachyarrhythmias self-terminated within seconds. Then, the next burst pacing was applied with at least 3 min of intervals. When tachyarrhythmia was sustained for more than 300 s, no further burst pacing was introduced. Once electrophysiology procedures were completed, the mice were euthanized under deep anesthesia by exsanguinations without recovery. Then, tissues were collected for further histological or expression studies.

### Isolated heart optical mapping

Mouse hearts were isolated and perfused retrogradely with 37 °C Krebs-Henseleit (KH, NaCl 118 mM, KCl 4.0 mM, KH_2_PO_4_ 1.2 mM, CaCl_2_ 1.8 mM, MgSO_4_ 1.6 mM, NaHCO_3_ 25 mM, glucose 5.5 mM, creatine 0.038 mM, and Na-pyruvate 2 mM) buffer saturated with 95% O_2_ and 5% CO_2_ at a constant flow rate of 2-3 mL/min to maintain baseline coronary pressure at 80-100 mmHg. The right atrium was electrically paced with rectangular pulses (2 ms width) at 7-8 Hz or burst paced (100 Hz x 10 s) using silver wire electrodes. Membrane potential dye, Di-4-ANEPPS (AnaSpec, Catalog# AS-84723, 1 µM in KH buffer), was perfused to image excitation of the heart. The dye was excited using a high-powered LED at 530 nm (LEX2-LG4, BrainVision, Tokyo, Japan). The emitted fluorescence (580 nm long pass) was detected using a high-speed CMOS camera (MiCAM03, BrainVision, Tokyo, Japan) with BV Workbench (version 1.13) acquisition software. BV-ana (version 16.04) and BV Workbench were used to analyze captured images to reconstruct movies, activation maps and phase maps. CCh was perfused at 1 µM in KH buffer.

### Isolated atria optical mapping

Both atria were isolated and placed in a superfusion chamber (Scientific Systems Design, Ontario, Canada). Atrial tissues were superfused with 37 °C KH buffer saturated with 95% O_2_ and 5% CO_2_ at a flow rate of 3 mL/min. Di-4-ANEPPS (2 µM in KH buffer) was superfused to image membrane potential-dependent signals. Stainless steel pin electrodes were used to deliver electrical stimulation to the RA appendage (RAA).

### Transverse aortic constriction

Male 10 week old C57BL/6J mice (The Jackson Laboratory) were subjected to TAC or sham surgery. Animals anesthetized with isoflurane were intubated and ventilated. After adequate depth of anesthesia, the chest cavity was opened by an incision of the left second intercostal space and the aortic arch was dissected from the surrounding tissues. The aortic arch was constricted against a 27 gauge needle with 7-0 silk suture. When the needle was removed, and the aortic constriction was generated with a cross-sectional area equivalent to the area of the 27 gauge needle. The chest cavity, muscles and skin were closed. In sham control mice, the entire procedure was identical except that the constriction of the aortic arch was omitted. Analgesic agents, meloxicam and buprenorphine were provided in each animal preemptively and post-operatively for 3 days. After the surgery animal will be monitored at least daily for pain, distress, sepsis and activity until the incision cite was healed, and sutures were removed. At 8, 12 and 16 weeks after surgery, groups of animals were subjected to terminal electrophysiology to evaluate AF inducibility as described above, and hearts were taken for histological evaluation.

### Angiotensin II infusion

Ten weeks old male C57BL/6J mice were subcutaneously implanted with minipumps (Alzet osmotic pump #1004). Preemptive and 3 days postoperative analgesia was provided to the mice. Ang II (Sigma-Aldrich, Catalog# A9525, 2 mg/kg/day, n = 8) or vehicle (0.01 N acetic acid in saline, n = 8) was infused continuously for 3 weeks. At 2 weeks after treatment initiation, cardiac function was evaluated using echocardiography. At 3 weeks after treatment initiation, animals were subjected to terminal electrophysiology to evaluate AF inducibility as described above and hearts were taken for histological evaluation.

### Echocardiography

Cardiac ultrasound images were obtained from isoflurane anesthetized mice using a Vevo 3100 system (FujiFilm VisualSonics, Ontario, Canada). Left atrial diameter was measured in the parasternal long axis view from the anterior to posterior walls. Left ventricle dimensions and systolic functions were quantified from images obtained in the parasternal short axis view obtained in M-Mode at the mid-level of the papillary muscles. Isovolumic relaxation time (IVRT) was evaluated using pulsed wave Doppler in the apical 4-chamber view. All images were analyzed using VevoLab software.

### Heart specific LKB1-deficient mice

LKB1 floxed mice were obtained from The Jackson Laboratory (Stock No: 014143) [27], as were αMHC-Cre transgenic mice (Stock No: 011038). The strains were interbred and LKB1^flox/flox^ αMHC-Cre^+/-^ mice were used as a cardiac-specific deficient strain (Cardiac-LKB1 KO); littermate LKB1^flox/flox^ αMHC-Cre^-/-^mice were used as controls. The genotypes of the mice were confirmed by PCR using protocols from The Jackson Laboratory. Cardiac-LKB1 KO and control mice were implanted with telemetry devices with ECG leads, which were subcutaneously secured at right shoulder and left flank to obtain ECG lead II (HD-X11, Data Sciences International, St. Paul, MN) at 5 weeks old. Preemptive and 3 days postoperative analgesia was provided to the telemetry implanted mice. Ambulatory ECG signals were acquired and analyzed for 3 days a week for 4 weeks immediately after telemetry implantation using Ponemah software (version 6.50, Data Sciences International, MN, USA). Once data acquisition was completed, the mice were euthanized by exsanguinations under deep anesthesia. Tissues were collected for further histological or expression studies.

### mRNA expression and histology analysis

Heart tissues were either snap frozen with liquid nitrogen for mRNA extraction or fixed with formalin for the histological examinations. Total RNA was extracted using RNeasy Fibrous Mini kit (QIAGEN, Cat# 74704). cDNA was synthesized using SuperScript VILO Master Mix (Thermo Fisher Scientific, Catalog# 11755250). mRNA levels were quantified using gene-specific primer pairs by real-time RT-PCR on an ABI PRISM 7900HT Sequence Detection System (Thermo Fisher Scientific, MA, US) according to the manufacturer’s protocol. Gene expression was normalized to the geometric mean of three internal control genes (Actin, β-2-microglobulin and TATA-box binding protein). Assay reagents for each gene were purchased from Thermo Fisher Scientific (Supplemental Table S1). For histological assessment of fibrosis, atria were separated from ventricles and embedded for a long-view (sagittal) orientation after marking the right atria with a blue tissue dye to aid in identification. Samples were microtome sectioned at 5 µm thickness, and the sections were stained with ready-to-use Sirius red dye (Rowley biochemical, Catalog# SO-674) according to manufacturer’s protocol. Fibrotic area was quantified from a single mid-level section from each animal using the area quantification algorithm from the HALO image analysis program (Indica Labs, Inc.).

### Data analyses

Data were compared with those observed in the corresponding control group using t-tests with Bonferroni correction (GraphPad Prism™ 8.03). Data are expressed as mean ± SEM. Arrhythmia inducibility was calculated as % of positive tachyarrhythmias from total burst pacing stimulations. Arrhythmia inducibility was compared with the corresponding control group using χ square test. Cumulative arrhythmia times were not distributed normally, and therefore the parameters was expressed as median ± interquartile range (IQR). Statistics were applied non-parametric comparisons with either Mann-Whitney U test or Kruskal-Wallis test.

## Results

### CCh and burst pacing-induced atrial tachyarrhythmia *in vivo* and in isolated hearts of mice

Acute CCh treatment decreased heart rate in anesthetized C57BL/6 mice (537 ± 5 vs. 360 ± 23 bpm, vehicle and CCh, respectively). Fig. 1A and 1B show representative ECG and RA EG traces of vehicle- and CCh-treated mice, respectively. Vehicle-treated animals (Fig. 1A) showed sinus rhythm immediately after burst pacing. In contrast, CCh treated mice (Fig. 1B) showed a high frequency activation pattern on RA EG and irregular RR interval on ECG. The right atrial ERP was significantly decreased in the CCh-treatment group (Fig. 1C). Burst pacing-induced tachyarrhythmia occurrence and cumulative tachyarrhythmia duration were significantly increased by CCh treatment (Fig. 1D and 1E, respectively). In isolated hearts, left atrial membrane potential was imaged using an optical mapping system. Despite an AF-like phenotype by ECG and atrial EG, the burst pacing-induced tachyarrhythmia in CCh mice showed very similar excitations and propagation patterns to those of sinus rhythm without any reentrant signals or wavelets in the left atrium (Fig. 1F and Supplemental Movie S1). The excitation was regular and high frequency (∼50 Hz, Fig. 1G), and therefore not a faithful representation of human AF. We observed similar regular high frequency excitation in all (n = 3) mouse left atria in isolated hearts. In contrast, spiral waves were detected during ventricular fibrillation in mouse isolated hearts using phase mapping (Supplemental Fig. S1.). The demonstration of the ability to detect ventricular reentry in this system indicates that the lack of atrial reentry was *bona fide* and not due to technical limitations.

**Fig. 1.**
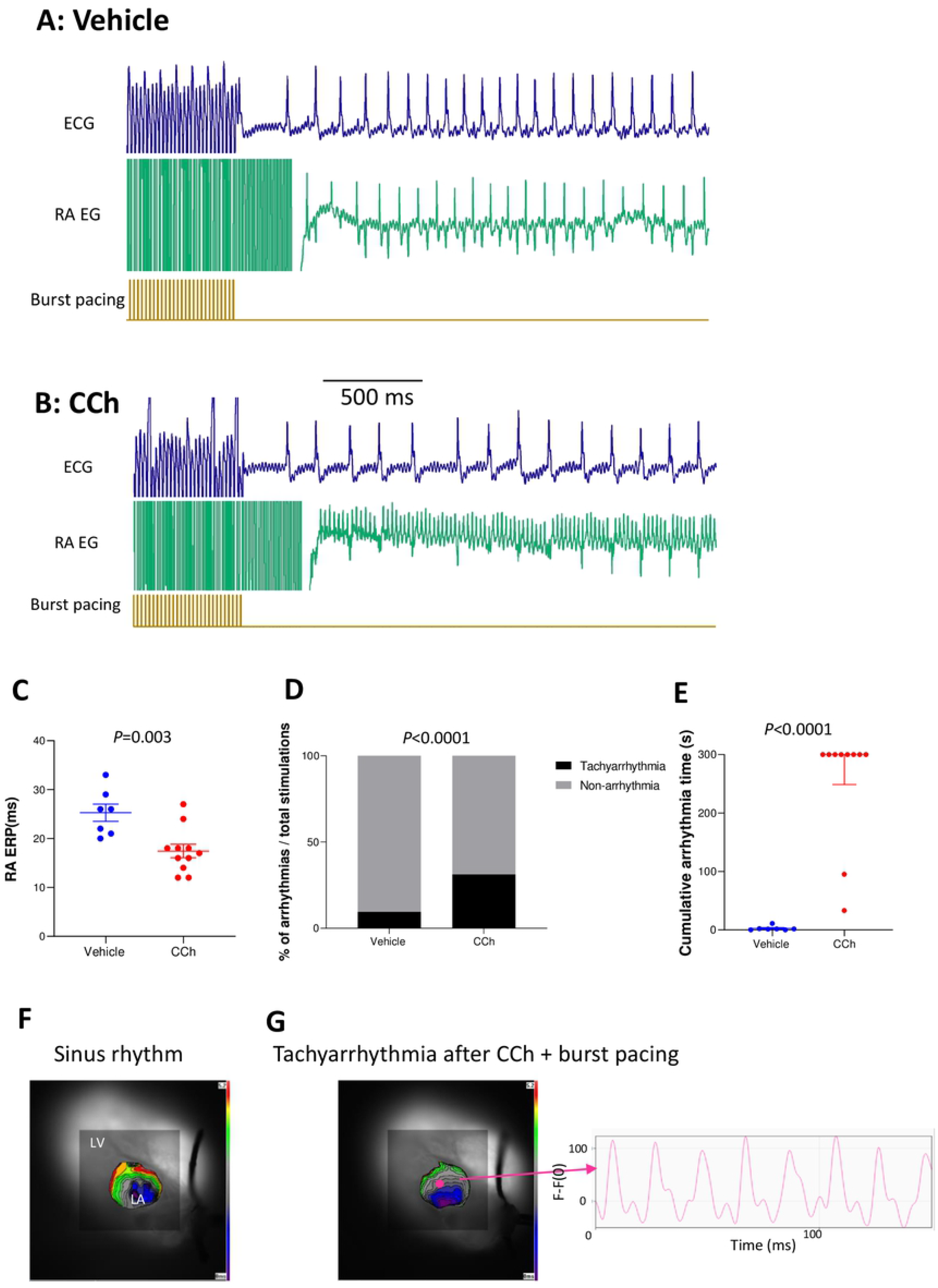
Characteristics of CCh + burst pacing-induced tachyarrhythmia in C57BL/6 mice and in isolated hearts. A and B: representative ECG and RA EG traces after 50 Hz burst pacing in vehicle-treated and CCh-treated (0.3 mg/kg, ip) mice, respectively. C: Right Atrial effective refractory period (RA ERP) in vehicle-treated and CCh-treated groups. Each point indicates individual animal data, and lines represent mean ± SEM. D: % of total burst pacing stimulations that resulted in atrial tachyarrhythmias in each group. E: Cumulative atrial tachyarrhythmia time in vehicle and CCh groups. Each point indicates individual animal data, and lines represent median ± IQR. F: Representative activation map during sinus rhythm before CCh treatment. LA activation starts from the lower right and propagates to the upper left. G: An activation map of the same heart after CCh and burst pacing. The LA shows an induced tachyarrhythmia with an activation pattern that is very similar to sinus rhythm, but with high and regular frequency (∼50 Hz). Similar observations were obtained from 3 out of 3 hearts.

### Atrial tachyarrhythmia vulnerability of TAC or Ang II infusion mice

The above observations from acute CCh-treated mice did not show convincing data representative of human AF. Next, we tested murine models of atrial structural remodeling in an effort to identify a model with more representative features of human AF. The TAC surgery-induced cardiac hypertrophy model was reported to be vulnerable to burst pacing-induced AF [11-14, 20]. In our studies, TAC surgery increased the heart weight to body weight ratio in C57BL/6 mice (Fig. 2A), reduced ventricular ejection fraction (Fig. 2B, 62 ± 2% and 52 ± 2%, sham and TAC respectively), and increased LA diameter (Fig. 2C, 2.6 ± 0.1 mm and 3.1 ± 0.1 mm, sham and TAC respectively) at 3 weeks post-surgery relative to sham-operated controls as determined by echocardiography. Those parameters exhibited trends in the same direction at 7 and 11 weeks after surgery. LA fibrotic area was increased significantly in TAC mice at 16 weeks (Fig. 2D, E, 12.6 ± 1.0% vs 16.4 ± 0.6%, sham and TAC respectively). There were no changes in RA ERP between sham and TAC mice at 8, 12 or 16 weeks post-surgery (Fig. 2F). Right atrial burst pacing-induced atrial tachyarrhythmia vulnerability was not different between sham and TAC mice at any time point (Fig. 2G, H). Thus, we observed no evidence of an AF-like phenotype in TAC mice. We further evaluated the severe pressure overload model of ascending aortic constriction (AAC). Heart size and RA ERP were significantly increased at 5 weeks after surgery in AAC group. Burst pacing-induced atrial tachyarrhythmia vulnerability was not different between sham and AAC groups (S2 Fig).

**Fig 2.**
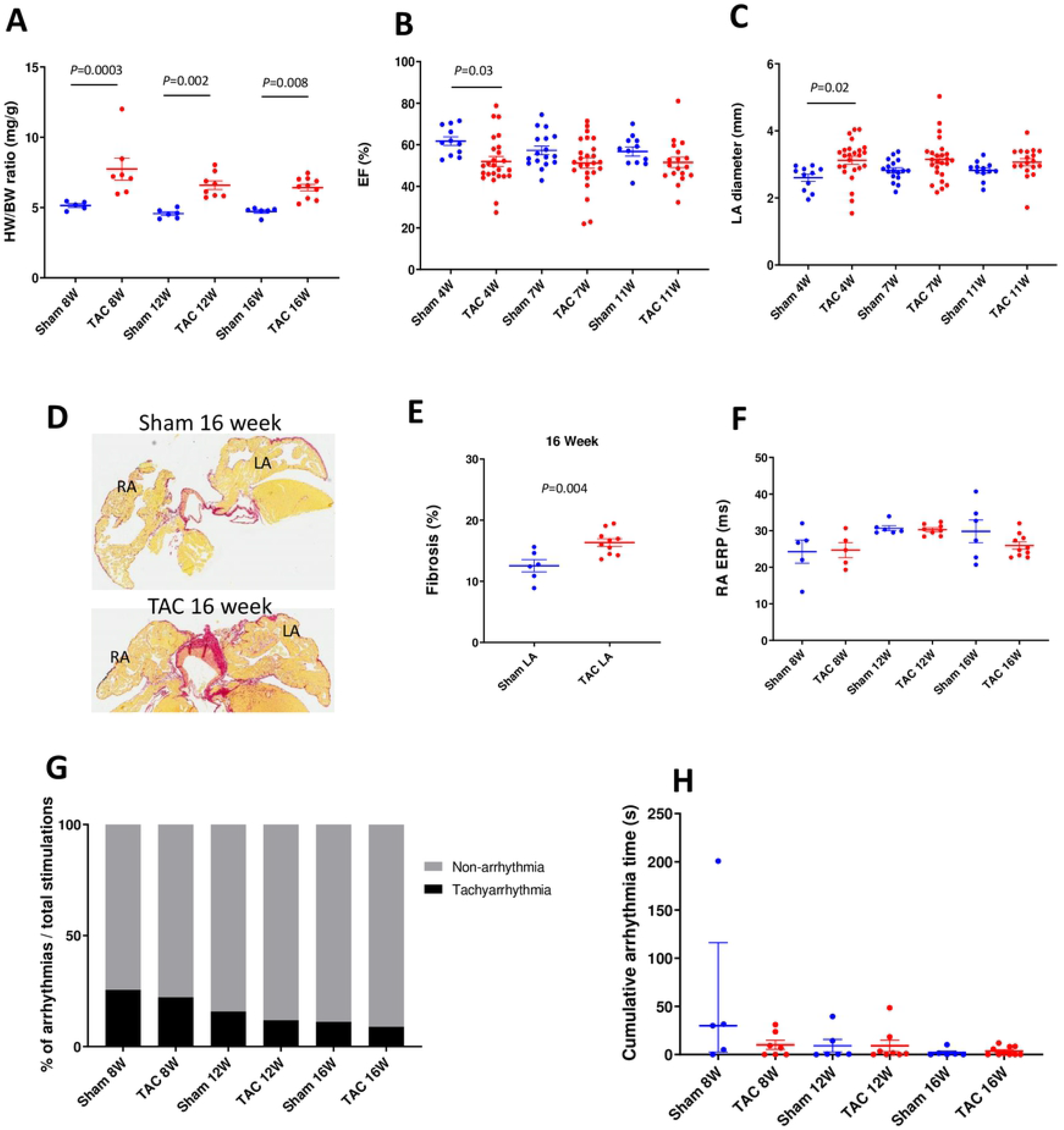
Heart phenotypes and burst pacing-induced tachyarrhythmia in TAC mice. A: Heart weight / body weight ratio in sham and TAC mice at 8, 12 and 16 weeks after surgery. Echocardiographic evaluation of ejection fraction (B) and LA diameter (C) in sham and TAC mice at 3, 7 and 11 weeks after surgery. D: Representative images of fibrosis staining (Picrosirius red) of the sham and TAC surgery mouse atria at 16 weeks after surgery. E: LA fibrotic area in sham and TAC mice at 16 weeks after surgery. F: RA ERP in sham and TAC mice at 8, 12 and 16 weeks after surgery. Each point indicates individual animal data, and lines represent mean ± SEM. G: % of positive atrial tachyarrhythmias from total burst pacing stimulations in sham and TAC groups at 8, 12 and 16 weeks after surgery. H: Cumulative atrial tachyarrhythmia time in sham and TAC groups. Each point indicates individual animal data, and lines represent median ± IQR.

Dosing Ang II via infusion has also been reported to induce cardiac structural remodeling with increased AF vulnerability [7-10, 28, 29]. Three weeks of Ang II infusion increased heart size in C57BL/6 mice (Fig. 3A), but did not change EF or LA diameter (Fig. 3B, C). Atrial fibrotic area was increased in left atria relative to vehicle controls (Fig. 3D, 9.7 ± 0.4% vs 18.3 ± 1.9%, vehicle and Ang II respectively). There were no changes in RA ERP between vehicle and Ang II infusion mice (Fig. 3F). Percent tachyarrhythmia occurrence or cumulative tachyarrhythmia time was not changed by Ang II infusion despite these structural changes (Fig. 3G, H). Thus, like the CCh-treated and TAC mice, we observed no evidence of an AF-like phenotype in Ang II infusion mice.

**Fig 3.**
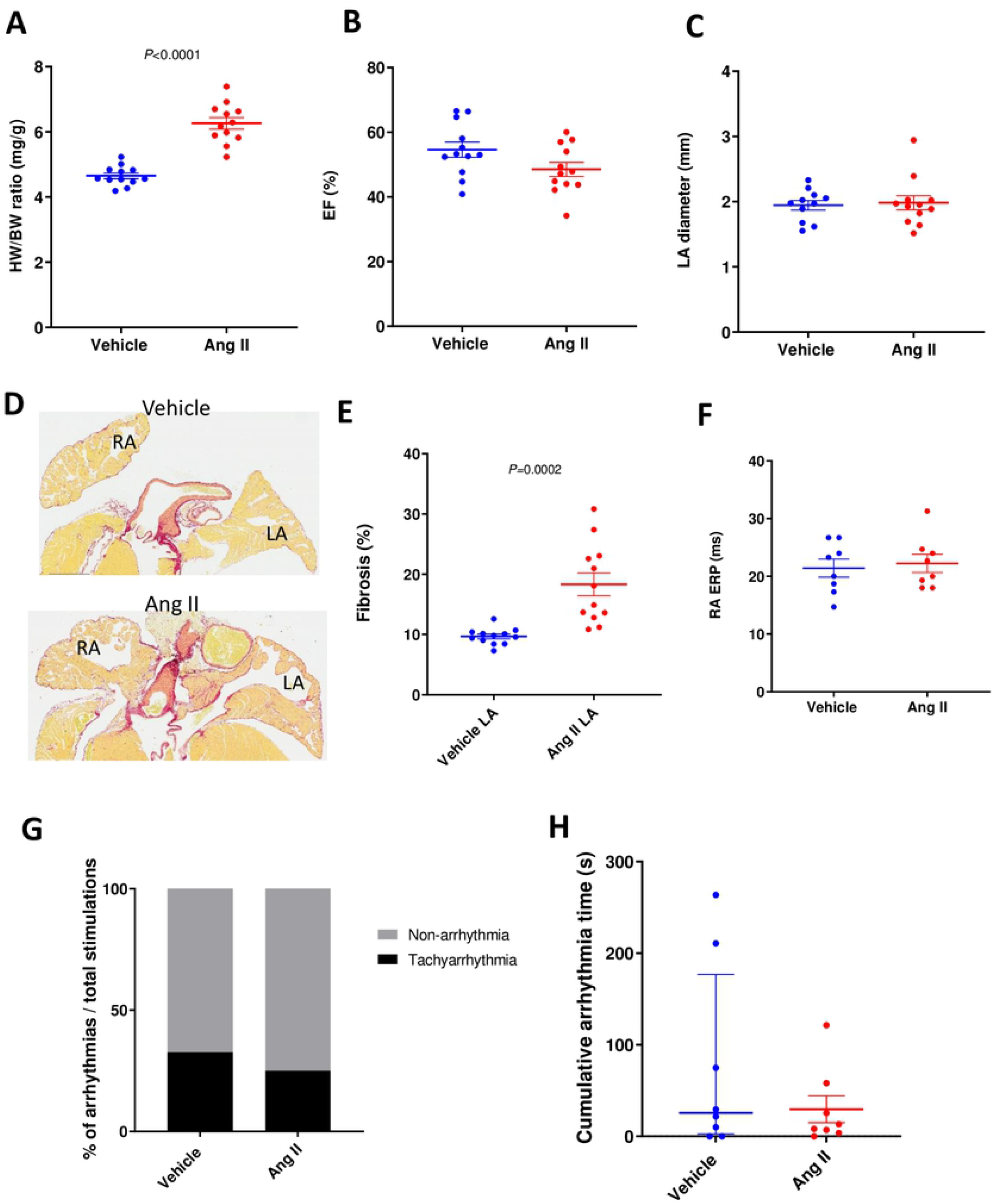
Heart phenotypes and burst pacing-induced tachyarrhythmia in Ang II mice. A: Heart weight / body weight ratio in vehicle and Ang II infusion mice. Echocardiographic evaluation of ejection fraction (B) and LA diameter (C) in vehicle and Ang II infusion mice. D: Representative images of fibrosis staining of the vehicle and Ang II infusion mouse atria. E: LA fibrotic area in vehicle and Ang II infusion mice. F: RA ERP in vehicle and Ang II infusion mice. Each point indicates individual animal data, and lines represent mean ± SEM. G: % of positive atrial tachyarrhythmias from total burst pacing stimulations in vehicle- and Ang II-treated groups. H: Cumulative atrial tachyarrhythmia time in vehicle and Ang II groups. Each point indicates individual animal data, and lines represent median ± IQR.

### ECG, heart phenotype and electrophysiology of Cardiac-LKB1 KO mice

Since we did not observe human-like AF in mouse models featuring burst pacing, we next profiled Cardiac LKB1 KO mice, which have been reported to have a spontaneous AF phenotype. Cardiac-LKB1 KO mice ECGs were monitored via telemetry. At 5 weeks of age, all cardiac-LKB1 KO mice (n = 8) showed an AF-like ECG phenotype with irregular RR intervals without discernable P waves (Fig. 4A), whereas all control mice (n = 6) showed normal sinus rhythm. Echocardiography revealed a significant increase in LA diameter (Table 1) in the Cardiac-LKB1 KO mice compared to control mice. Cardiac-LKB1 KO mice had enlarged RA and LA (Fig. 4B), and more than half of the Cardiac-LKB1 KO mice had thrombi in the LA appendage (LAA), observed post mortem (5 out of 8 in Cardiac-LKB1 KO mice vs. 0 out of 8 in control mice). Histological evaluation revealed increased fibrotic area in LA in Cardiac-LKB1 KO mice (18.0 ± 1.4 vs 38.5 ± 1.8%, Fig. 4B and C). In contrast, ventricular functions including EF, stroke volume and left ventricular (LV) mass were not different between control and Cardiac-LKB1 KO mice (Table 1). Left ventricular isovolumic relaxation time indexed to RR interval (IVRT/RR) was significantly increased in Cardiac-LKB1 KO mice.

**Fig. 4.**
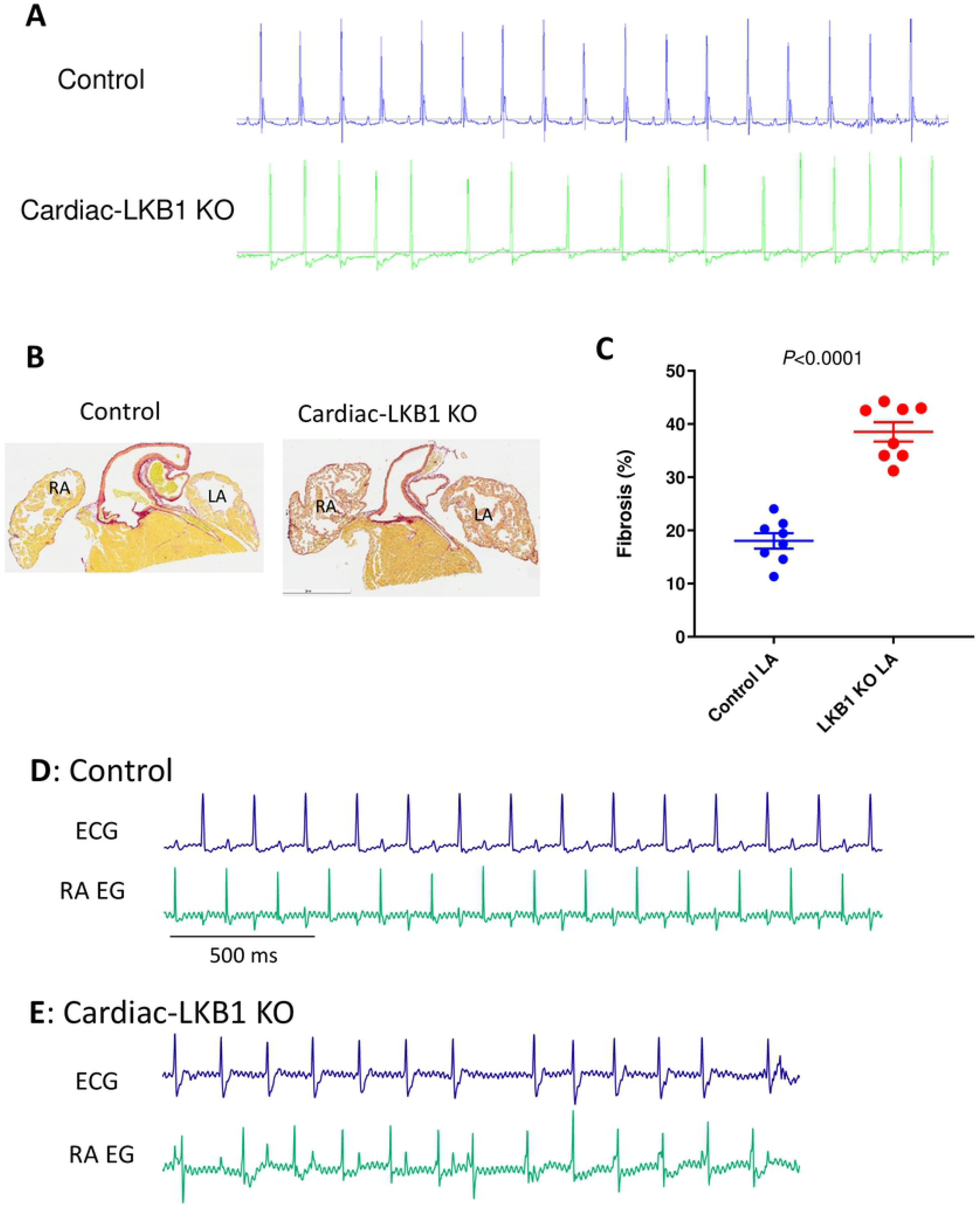
ECG, heart histology and RA electrogram traces in Cardiac-LKB1 KO mice. A: Representative ECG traces from telemetry-implanted control and Cardiac-LKB1 KO mice. B: Representative images of fibrosis staining of the control and Cardiac-LKB KO mouse atria. Cardiac-LKB1KO mouse had increased red staining, indicative of collagen positive areas, in the atria relative to control atria. C: % of positive red stain area over entire LA and RA tissue in control and Cardiac-LKB1 KO mice. Each point indicates individual animal data, and lines represent mean ± SEM. D: Control mice showed sinus rhythm, clear P waves and corresponding RA signal on RA EGs under anesthesia. E: Cardiac-LKB1 KO mice showed irregular RR intervals without P waves on ECG. RA EGs revealed independent and irregular atrial signals.

**Table 1.** Echocardiographic parameters of control and Cardiac-LKB1 KO mice.

|  | Control | Cardiac-LKB1 KO |
| --- | --- | --- |
| EF (%) | 59.7 ± 1.4 | 54.8 ± 3.5 |
| LVIDd (mm) | 3.81 ± 0.03 | 3.79 ± 0.10 |
| Stroke volume (μL) | 37.4 ± 1.3 | 33.8 ± 2.5 |
| LA diameter (mm) | 2.15 ± 0.06 | 3.23 ± 0.07*** |
| LV mass (mg) | 68.6 ± 2.8 | 69.8 ± 3.1 |
| IVRT/RR | 0.130 ± 0.002 | 0.23 ± 0.18*** |

LA, left atrium; LV, left ventricle; LVIDd, diastolic LV-internal diameter, IVRT/RR, isovolumic relaxation time indexed to RR interval. N = 10-12. Values are expressed as mean ± SEM.
\*\*\* $P < 0.001$ .

To examine excitation of the atria, RA EG were compared to corresponding surface ECGs, measured concurrently in mice under anesthesia. Control mice (n = 7) had sinus rhythm and, as expected, atrial excitations observed in the EGs aligned with the P waves measured on ECGs (Fig. 4D). In contrast, while Cardiac-LKB1 KO mice did exhibit apparent atrial excitation, these were irregular in nature, and were never observed to have a high-frequency, fibrillatory pattern (7 out of 7 mice). Furthermore, there was no discernable relationship between these atrial signals and ventricular depolarizations observed via ECG, indicating that the ventricular excitations were not governed by atrial excitations (Fig. 4E).

To further profile atrial excitation, Cardiac-LKB1 KO mouse isolated intact hearts and isolated atria were examined via optical mapping. Sinus rhythm and atrial excitation were detected in perfused control mouse hearts (Fig. 5A and Supplemental Movie S2) but were never detected in the right or left atria in Cardiac-LKB1 KO mice (Fig. 5B and Supplemental Movie S3).

**Fig. 5.**
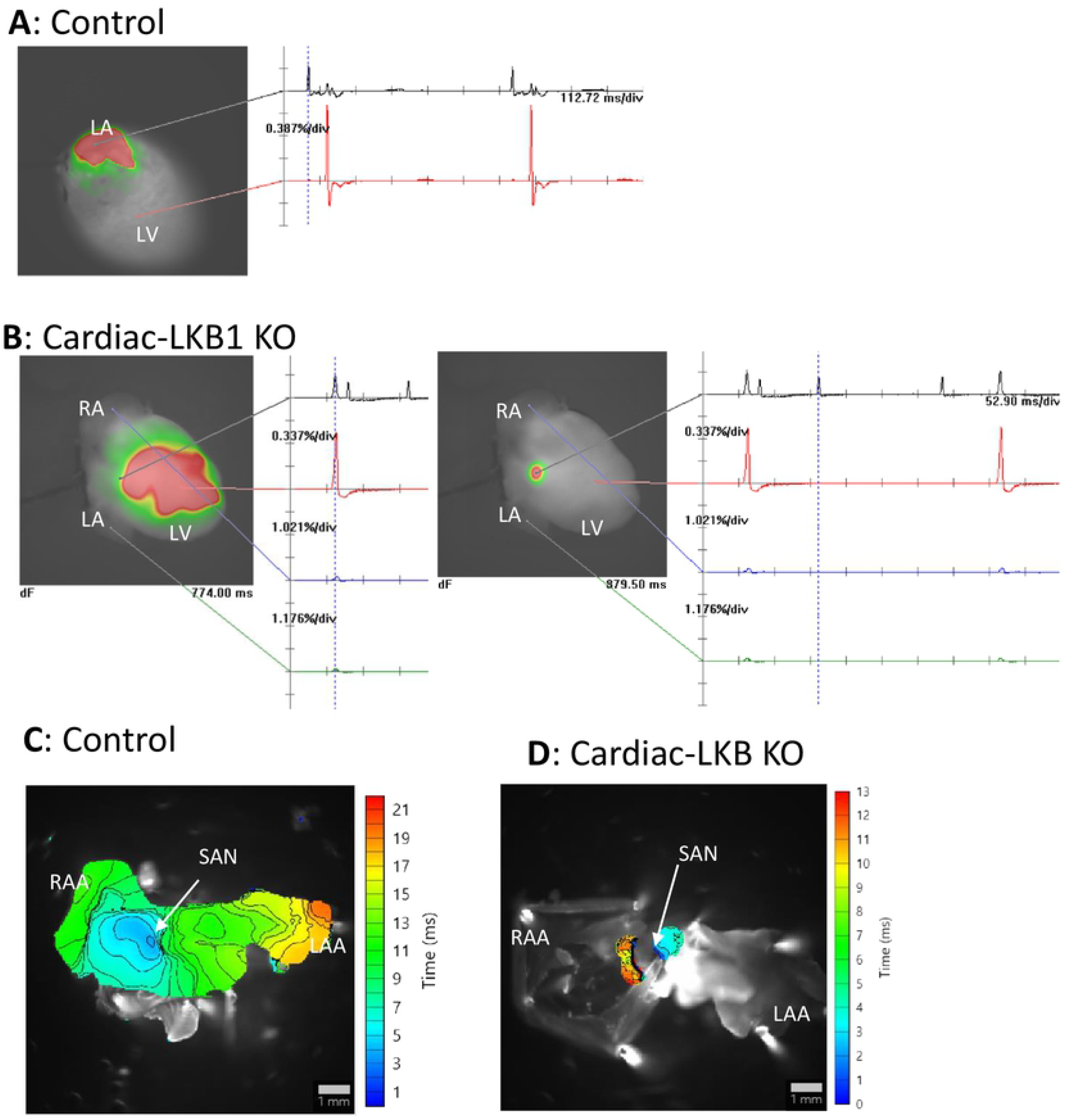
Optical mapping of control and Cardiac-LKB1 KO mouse hearts. A: Representative image of a perfused control mouse heart. LA excitation was detected, and the heart was in sinus rhythm. B: Representative images of perfused Cardiac-LKB1 KO mouse hearts. No atrial excitation was detected and ventricular excitation was confirmed (left). The right image shows a focal area close to atrial septum that was excited independently and irregularly. C: Representative activation map of control mouse atria. Excitation was initiated from the SAN, then propagated though the RA and LA and ended in the RAA and LAA. D: A representative activation map of Cardiac-LKB1 KO mice atria. Excitation was observed only at SAN area; the excitation did not propagate through the atria.

Interestingly, independent excitation generation sites were detected in the RA close to the intra-atrial septum (6 out of 7, Fig. 5B, Supplemental Movie S3 and S4) in Cardiac-LKB1 KO mice, but these excitations did not propagate to the whole atrium. These patterns of focal excitation were consistent with those of RA EGs in anesthetized Cardiac-LKB1 KO mice. We next used isolated and superfused atrial tissue to evaluate precisely the site of the focal excitations. In control mouse atria, excitations propagated from the sino-atrial node (SAN) to the RAA as well as to the LAA (Fig. 5C and Supplemental Movie S5). In contrast, in Cardiac-LKB KO mice, only the SAN area showed excitations (Fig. 5D and Supplemental Movie S6); the excitation was not propagated throughout the atria. This lack of atrial excitation was also evident when exogenous electrical stimulation was applied to the RAA (Supplemental Movie S7, 6 out of 6 preparations). We also profiled RA EGs of heterozygous Cardiac-LKB1 KO (LKB1^flox/-^αMHC-Cre^+/-^) mice. The heterozygous Cardiac-LKB1 KO mice did not show any of above phenotypes (Supplemental Fig S3). Therefore, severe cardiomyocyte knock down of LKB1 is likely required to show these pathological phenotypes.

### Gene expression of the right and left atria and left ventricle in Cardiac-LKB1 KO hearts

Expression of key genes was evaluated via RT-PCR in control and Cardiac-LKB1 KO hearts. LKB1 expression was reduced by more than half in both the atria and left ventricle of Cardiac-LKB1 KO mice (Fig. 6). Expression of the fibrosis related gene transforming growth factor β1 (TGFb1) was decreased in Cardiac-LKB1 KO mouse atria compared to control mouse atria, whereas other fibrosis related genes, collagen type 1 α1 (COL1a1) and connective tissue growth factor (CTGF), were increased. COL1a1 expression was also increased in Cardiac-LKB1 KO mouse ventricle. Strikingly, there was almost no SCN5a expression in Cardiac-LKB1 KO mouse atria. Connexin 40 (CX40) expression was robust in the atria but not in the left ventricle of control mice, whereas in Cardiac-LKB1 KO mouse hearts, very little CX40 expression was detected. This lack of SCN5A and CX40 expression suggests a profound conduction deficit in the LKB1 KO mice atria.

**Fig 6.**
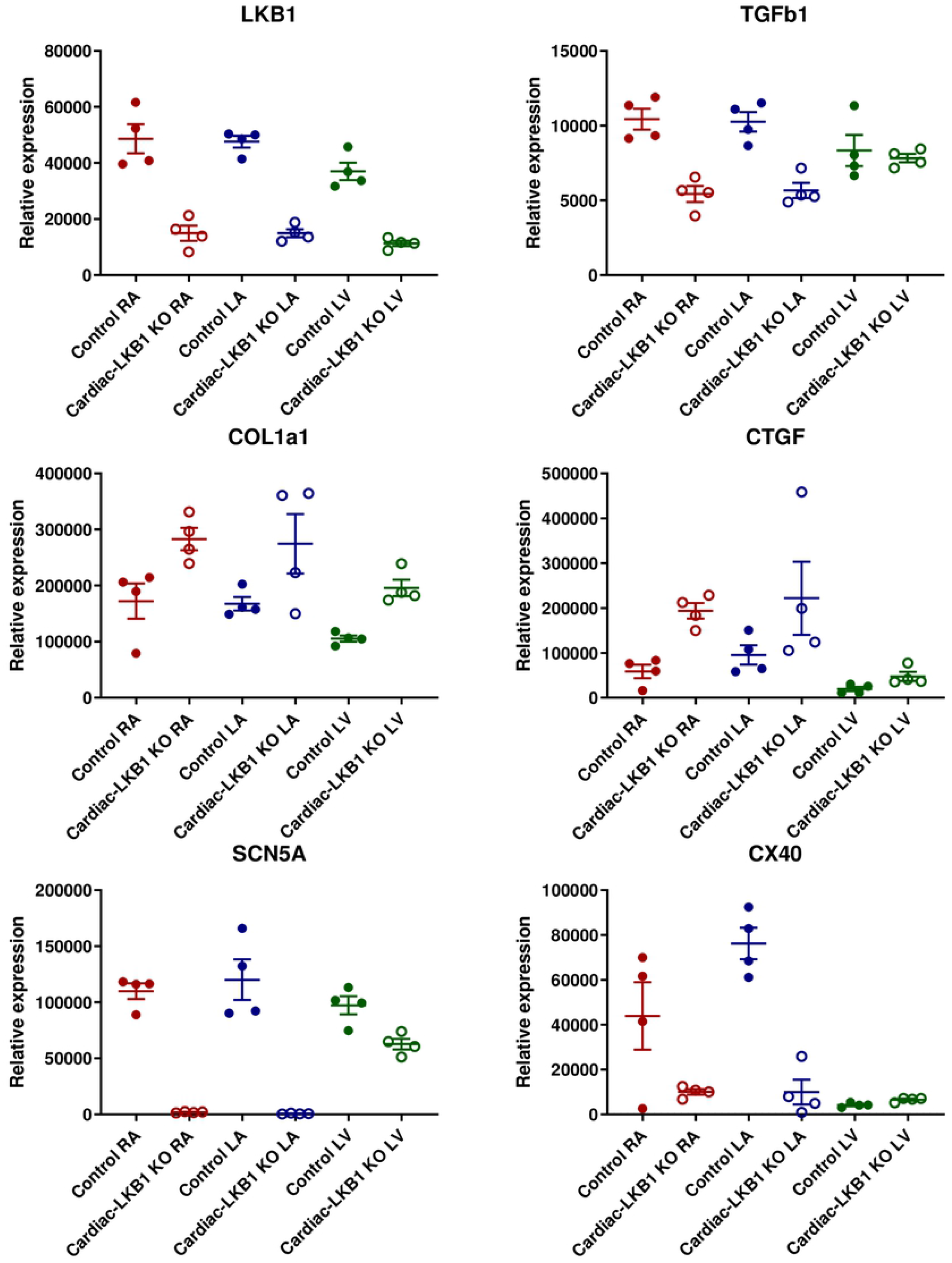
mRNA expression level in right atria, left atria and left ventricles of Cardiac-LKB1 KO and control mice. mRNA expression quantification for LKB1, TGFβ1 (transforming growth factor β1), COL1a1 (collagen type 1 α1), CTGF (connective tissue growth factor), SCN5a (voltage gated sodium channel α subunit 5) and CX40 (connexin 40). Each point indicates individual animal data, and lines represent mean ± SEM.

## Discussion

In order to identify a mouse model that mimics human AF pathophysiology, we first studied an acute model: CCh treatment with burst pacing. The rationale behind this model is that cholinergic activation induces GIRK channel opening to increase *I*KACh [30, 31], thereby reducing atrial action potential duration and ERP to promote reentry. In fact, GIRK is known to be activated in human AF as a part of ion channel remodeling [30, 32]. Indeed, we found that acute treatment with CCh decreased atrial ERP and increased burst pacing-induced vulnerability to atrial tachyarrhythmia in anesthetized mice. However, when we evaluated CCh and burst pacing-induced tachyarrhythmia using optical mapping in isolated perfused hearts, the atrial tachyarrhythmias in the left atrium were not typical AF-like reentrant arrhythmias; rather they resembled high and regular frequency cycles of atrial flutter-like arrhythmia. Atrial flutter is known to feature macro-reentry [33]. Limitations of this study are that 1) we could not capture the whole right and left atria simultaneously, due to challenges to avoid artifacts from complexed 3D surface structure of atria [34, 35] and 2) *in vitro* isolated heart characteristics and *in vivo* situation might differ. Therefore, we might have missed macro-reentry across both atria or micro-reentry in *in vivo*. Alternatively, automaticity of the SAN or other triggered activities may induce such high frequency firing [36]. While we did not observe micro-reentry in the mouse atria in isolated hearts, we did detect it in the ventricles. Therefore, it is unlikely we missed micro-reentry of the atria for technical reasons.

We next evaluated the widely used murine heart disease models of TAC surgery or Ang II infusion, which have been reported to increase AF vulnerability [7-14]. These models are known to induce structural and electrical remodeling in both atria and ventricles, which is not a feature of the acute CCh model. Structural remodeling such as fibrosis can be a substrate for micro-reentrant circuits during AF. In the TAC mice, HW/BW ratio, LA diameter and LA fibrosis were increased. However, we were not able to demonstrate increased atrial tachyarrhythmia vulnerability, nor did we observe changes in atrial ERP in the TAC mice. Some TAC surgery model studies reported both increased atrial fibrosis and AF vulnerability [11, 13, 14], with more than a 3-fold increase of atrial fibrosis. In contrast, our data indicated ∼30% increases in fibrosis in TAC mice. We therefore thought that our TAC surgery model might not been sufficiently severe and/or lacked substrates for AF, and evaluated a more severe model of TAC, the ascending aortic constriction model. However, again we did not increase AF vulnerability.

In contrast to pressure overload models, increased fibrosis was more evident in left atria in the Ang II-treated mouse model. Increased heart weight was also noted in Ang II-treated mice. However, once again we were not able to demonstrate increased atrial tachyarrhythmia vulnerability, nor did we observe changes in atrial ERP in the Ang II mice. Heterogeneity of the local refractoriness or action potential duration is also a key factor for development of reentrant circuits [37]. These properties may have been missing in our TAC and Ang II models despite the increased atrial fibrosis.

Due to the difficulty in validating an inducible AF model in our laboratory, we evaluated Cardiac-LKB1 KO mice, which were reported to have spontaneous AF featuring many similar pathologies to human AF [23-25]. The LKB1-AMPK pathway is involved in the remodeling pathway for ion channels, fibrosis and apoptosis [38]. Consistent with this, we observed increased atrial volume, severe atrial fibrosis and an irregular RR interval without P waves on ECGs. We also observed changes in atrial fibrosis-related genes such as decreased TGFβ1 and increased collagen in Cardiac-LKB1 KO mouse atria, the net effects of which may contribute to the observed atrial fibrosis. However, in contrast to the expected result of a *bona fide* AF-phenotype, we did not see any atrial fibrillation using either intra-atrial EGs or optical mapping. Atria in Cardiac-LKB1 KO mice were non-excitable even in the setting of exogenous electrical stimulations, and so we could not measure the atrial effective refractory period. The only atrial excitation observed were irregular, focal excitations at the SAN area, which did not convey through the whole atria. These isolated excitations seemed to contribute to the random intra-atrial EG signals. Therefore, although the Cardiac-LKB1 KO surface ECG phenotype is superficially reminiscent of typical AF, the mice actually have atrial standstill and/or myopathy. The ECG pattern observed was actually an escape rhythm. This may explain the high prevalence of thrombi deposition in LAA. The severely reduced expression of SCN5a and CX40 likely contribute to this non-excitable phenotype and extreme electrical remodeling. SCN5a normally initiates the atrial action potential. CX40 is known to be expressed only in atria, controlling conduction across atrial myocytes, and CX40 mutations are associated with AF [39-45]. Due to atrial standstill, Cardiac-LKB1 KO mice do not have the booster pump function of the left atrium, which may contribute to the increased IVRT [46]. Our Cardiac-LKB1 KO data suggest that a very careful phenotypic analysis is required to understand excitability defects in spontaneous murine AF models. ECGs, by themselves are not sufficient for this purpose. Rather than an AF model, the Cardiac-LKB1 KO model may represent an advanced atrial remodeling or myopathy model.

The hearts of mice, large animals and human have many similarities and differences in terms of structure, electrophysiology and size [47]. Reentrant circuits to sustain AF were confirmed in human [51, 52]. Large animals are usually taken as the “gold standard” AF models. For example, in dog atria, acute GIRK activation followed by burst pacing induced micro-reentrant circuits with dispersed dominant frequency [48-50]. In contrast, rodent models of AF have been arguably less reliable. For example, De Jong et al. [53] reported that TAC-treated mice were not a suitable AF model, possibly due to insufficient substrates for AF. One of the long-standing hypothesis for lack of sustained reentry in mouse atria is insufficient critical mass of the mouse hearts [54]. Garrey [55] hypothesized almost 100 years ago that a certain size of myocardial tissue was required to sustain reentrant arrhythmias. Although Vaidya et al. [56] and we showed reentry or spiral wave in mouse ventricles, mouse atria are considerably smaller than ventricles. In addition, transmural persistent reentry is one of the drivers for AF in humans [57]. Mouse atria are extremely thin tissues that may not support transmural reentry. These size limitations may minimize occurrence of reentrant circuits in mouse atria. However, it is worth noting that others have reported reentrant type AF in mice using optical mapping imaging techniques similar to those employed in this study [19, 36, 58, 59]. One such model employed a sodium channel mutant [19]. In the model, increased fibrosis is reported in the atria. This could be a substrate for the reentrant circuit or spiral wave activity for arrhythmia in mouse atria. Therefore, atrial tachyarrhythmia and reentry patterns could be totally different from model to model.

In this study, we looked at four commonly used murine AF models, and found human-like atrial fibrillation in none of them. The data shown in this study suggest that to conclude any particular murine model is truly similar to human AF, careful examination, including use of atrial EGs and optical mapping, is required. It may be that other mouse models feature *bona fide* arrhythmia, but our own studies have led us to conclude that large animal models are the best, and possibly only, way to evaluate efficacy of anti-AF drugs intended for humans.

## Author contributions

All authors reviewed the manuscript; W.L., S.S.S., P.K.G., G.T., A.K.P.T., M.G.W., S.C., Y.A. formulated ideas, designed the study; F.F., M.P., L.C, S.P., O.T., J.L., J.M.L., Y.M., G.D., Y.A. performed experiments; F.F., S.P., W.L., J.L., J.M.L., Y.M., P.K.G., G.D., G.T., Y.A. analyzed experimental data; and Y.A. wrote the manuscript.

## Acknowledgments

Authors thank Lauren Janes and John Halupowski for their contributions to maintain and provide Cardiac-LKB1 KO mice.

## Supplemental information

**Supplemental Table S1. Reagents information for qRT-PCR**. Reagents information for mRNA quantifications in the evaluated genes and internal control genes. Since Thermo Fisher Scientific does not disclose primer sequences, assay identification number of each gene are shown.

**Supplemental Movie S1. Representative video of LA tachyarrhythmia in CCh + burst pacing isolated heart**. High frequent normal excitations are detected during tachyarrhythmia.

**Supplemental Movie S2. Representative video of perfused control mouse heart**. Heart is sinus rhythm. Left atrium following left ventricle excitations are detected.

**Supplemental Movie S3. Representative video of perfused Cardiac-LKB1 KO mouse heart**. Ventricular excitation is detected. While, there are no excitations in either right or left atria. There are focal and independent excitations in RA close to the atrial septum during ventricular excitations.

**Supplemental Movie S4. Representative video of perfused Cardiac-LKB1 KO mouse heart**. Ventricular excitation is detected. There are focal and independent excitations in RA close to the atrial septum during ventricular excitations.

**Supplemental Movie S5. Representative video of atrial preparation in control mouse**. Excitation is initiated from SAN. The excitation propagates to RAA and LAA

**Supplemental Movie S6. Representative video of atrial preparation in Cardiac-LKB1 KO mouse**. Excitation is limited in SAN area. The excitations do not propagate beyond RA

**Supplemental Movie S7. Representative video of atrial preparation in Cardiac-LKB1 KO mouse**. RAA is electrically stimulated. However, pacing excitations are not propagated. Independent SAN excitations are detected.

**Supplemental Fig. S1. The Cardiac-LKB1 KO heart with ventricular fibrillation**. Phase maps of the ventricular having ventricular fibrillation (VF). ∼30 Hz of spiral wave was observed. VF was established by several attempts of right ventricular 100 Hz burst pacing stimulations for 5 sec.

**Supplemental Fig. S2. Heart profiling and burst pacing induced tachyarrhythmias in AAC mice**. A: Heart weight / body weight ratio in sham and AAC mice at 5 weeks after surgery. B: Right atrial effective refractory period (ERP) in sham and AAC mice. Each point indicates individual animal data, and lines represent mean ± SEM. C: % of positive atrial tachyarrhythmias from total burst pacing stimulations in each group. D: Cumulative tachyarrhythmia time in each animal. Each point indicates individual animal data, and lines represent median ± IQR.

**Supplemental Fig. S3. Representative ECG and RA EG. Burst pacing induced tachyarrhythmias in Cardiac-LKB1 heterozygous KO mice**. A: Control mouse shows sinus rhythm, clear P wave and corresponding RA (RA) signal on RA EG. B: Cardiac-LKB1 heterozygous KO mouse also shows sinus rhythm. C: % of positive atrial tachyarrhythmias from total burst pacing stimulations in each group. D: Cumulative tachyarrhythmia time in each animal. Each point indicates individual animal data, and lines represent median ± IQR.

